# LRRK2 regulates the activation of the unfolded protein response and antigen presentation in macrophages during inflammation

**DOI:** 10.1101/2023.06.14.545012

**Authors:** Ahmed M. Fahmy, Tyler Cannon, Camberly Herdandez Paredes, Ali Ahmadi, Yong Zhong Xu, Benoit Barette, Maha Ibrahim, Joël Lanoix, Guillermo Arango-Duque, Erwin Schurr, Eric Chevet, Pierre Thibault, Samantha Gruenheid, Heidi McBride, Michel Desjardins

## Abstract

While the contribution of inflammation in the pathological process leading to Parkinson’s disease (PD) is well established, a growing body of evidence supports a role for the long-lasting adaptive immune system in the disease. We showed that, in inflammatory conditions, the PD-proteins PINK1 and Parkin negatively regulate the presentation of mitochondrial antigens on MHC I molecules, a process referred to as MitAP (**Mit**ochondrial **A**ntigen **P**resentation). In the absence of PINK1, over-activation of this pathway in antigen-presenting cells (APCs) engages autoimmune mechanisms leading to the establishment of cytotoxic CD8^+^ T cells. Co-culture of dopaminergic neurons (DNs) with these T cells led to neuronal cell death, suggesting that the MitAP in DNs made them susceptible to T cell-mediated destruction. In the present study, we used a pharmacological and genetic approach to characterize the MitAP pathway at the molecular level. We showed that this antigen presentation pathway is induced in APCs in response to inflammatory signals through the sequential activation of TLR4, cGAS-STING and the Unfolded Protein Response (UPR). A “UPR motif” present on a STING cytoplasmic domain was shown to specifically activate the UPR sensor IRE1. Remarkably, the PD-related protein LRRK2, acted with STING upstream of the UPR to regulate the transition from innate to adaptive immunity, thereby identifying this PD-related protein as a key player in the immune response during inflammation.

## Introduction

Parkinson’s disease (PD) is a progressive disorder that affects dopaminergic neurons (DNs) in the substantia nigra of the brain and causes movement impairment. While the neurodegenerative nature of PD is well established, the molecular mechanisms responsible for the initiation of the disease are still largely unknown, limiting the development of effective early stages therapeutic approaches. A large number of genes and genetic loci have been identified as risk factors for PD so far^1^, suggesting that complex and broad molecular pathways are likely to be at play in the disease process. Together with ageing, inflammation, a rapid innate immune response triggered during infection or tissue damage, has been identified as a key determinant for PD ^2^, pointing to a role of the immune system in PD ^3^. Post-mortem analyses have shown that T lymphocytes (T cells) are present in the brain of people with PD and animal models of the disease ^4,5^. This suggested that adaptive immunity, the long-lasting arm of the immune system engaged by the recognition of specific antigens by B and T cells, contributes to PD. In that context, the detection of autoreactive CD4^+^ T cells against α-synuclein peptides in people with PD emphasized the likeliness that autoimmune mechanisms play a role in PD pathophysiology ^6^.

We showed that two PD-related proteins, PINK1 and Parkin regulate the presentation of self-antigens of mitochondrial origin (MitAP) on MHC class I molecules at the surface of immune cells during inflammation ^7^. Indeed, we observed in cultured macrophages either expressing low levels or lacking PINK1 that infection with Gram-negative bacteria, as well as exposure to soluble lipopolysaccharide (LPS), significantly enhanced MitAP. Furthermore, we showed *in vivo* that gut infection with the Gram-negative bacterium *Citrobacter rodentium* in *Pink1*^*-/-*^ mice activated MitAP in antigen-presenting cells (APCs), leading to the establishment of a population of autoreactive cytotoxic CD8^+^ T cells. Remarkably, these mice which, otherwise, display a phenotype indistinguishable from their wild-type littermates, developed severe motor impairment within a few weeks, reversible by L-DOPA treatment ^8^. Since mitochondria-specific cytotoxic CD8^+^ T cells were observed in the central nervous system of infected *Pink1*^*-/-*^ mice, we hypothesized that T cells could recognize and attack DNs if these were to engage MitAP and present the corresponding mitochondrial antigens at their surface. While this remains to be confirmed *in vivo*, we observed that MitAP was induced in DNs from *Pink1*^*-/-*^ mice exposed to LPS and interferon-γ in culture ^8^. Co-culture of these neurons with a mitochondria-specific cytotoxic CD8+ T cell hybridoma in these conditions led to T cell activation and DNs attack.

These data suggested that two distinct instances of MitAP are required to activate a PD-like pathological process in this gut infection model. First, MitAP is triggered in APCs thus activating T cells in the periphery; second, MitAP is activated later in the disease process in DNs in the brain, rendering them susceptible to T cell attack. Thus, identifying key players involved in MitAP will provide insights for the development of therapeutic approaches for PD treatment.

In the present study, we used both pharmacological and genetic approaches to characterize how the MitAP pathway is activated and regulated in macrophages. Our data indicate that multiple stress sensors cooperate to induce MitAP. This integrated mechanism allows the transition from a transient innate inflammatory response to a long-lasting adaptive response driven by antigen presentation. Interestingly, this immune transition pathway requires the PD-related protein LRRK2 which acts in close relationship with STING, upstream of the activation of the ER stress sensor IRE1. The identification of key players in the MitAP pathway provides attractive targets for the development of therapeutic approaches aimed at both early and late stages of the PD disease process.

## Results

### LPS activates MitAP through TLR4

The binding of LPS, a pathogen-associated molecular pattern (PAMP) molecule present on the outer leaflet of the outer membrane of Gram-negative bacteria, to the Toll-like receptor 4 (TLR4) engages a complex inflammatory response ^9^. The finding that PINK1 and Parkin regulate the activation of MitAP in response to LPS suggested that these two PD-related proteins act along the TLR4 signaling pathway to modulate the engagement of the adaptive immune response during inflammation ^7^. To characterize how MitAP is triggered by LPS, we first compared the extent to which two Gram-negative bacteria, enteropathogenic *E. coli* (EPEC) and *Helicobacter pylori* (*H. pylori*), activate this pathway. We observed that while EPEC effectively activated MitAP, *H. pylori* was unable to do so (Fig. 1A). Similar results were observed with heat-killed bacteria (Fig. 1B), ruling out the possibility that MitAP activation required an active microbial process such as the release of microbial effectors to the cytoplasm during infection. These data go along with the finding that *H. pylori’*s LPS is unable to activate TLR4 signaling ^10^. We then determined whether MitAP could be induced by other microbial surface molecules. In addition to LPS, peptidoglycan and flagellin (recognized by TLR2 and TLR5 respectively) also activated MitAP, albeit at a much higher concentration (Figs. 1C-E). When cells were incubated with TAK242, a specific TLR4 inhibitor, MitAP activation by LPS was completely inhibited (Fig. 1F), confirming the involvement of this receptor in the presentation of mitochondrial antigens. TAK242 had no inhibitory effect when MitAP was induced by a high concentration of peptidoglycan and flagellin (Supp. Fig. 1). This ruled out the possibility that these molecule preparations were contaminated by LPS and rather suggested that TLR2 and TLR5 are much less effective at inducing MitAP than TLR4. All of these receptors share a common Myd88-dependent signaling arm, leading to the activation of NF-κB and the expression of pro-inflammatory cytokines such as IL-6^11^. Unlike TLR2 and TLR5, TLR4 engages a second signaling pathway involving the adaptor proteins TRAM and TRIF. This pathway activates the transcription factor interferon regulatory factor 3 (IRF3) and the expression of type-I interferons. The high potency of LPS suggested that the TRAM/TRIF of TLR4 signaling is involved in the activation of the MitAP pathway.

**Fig. 1:**
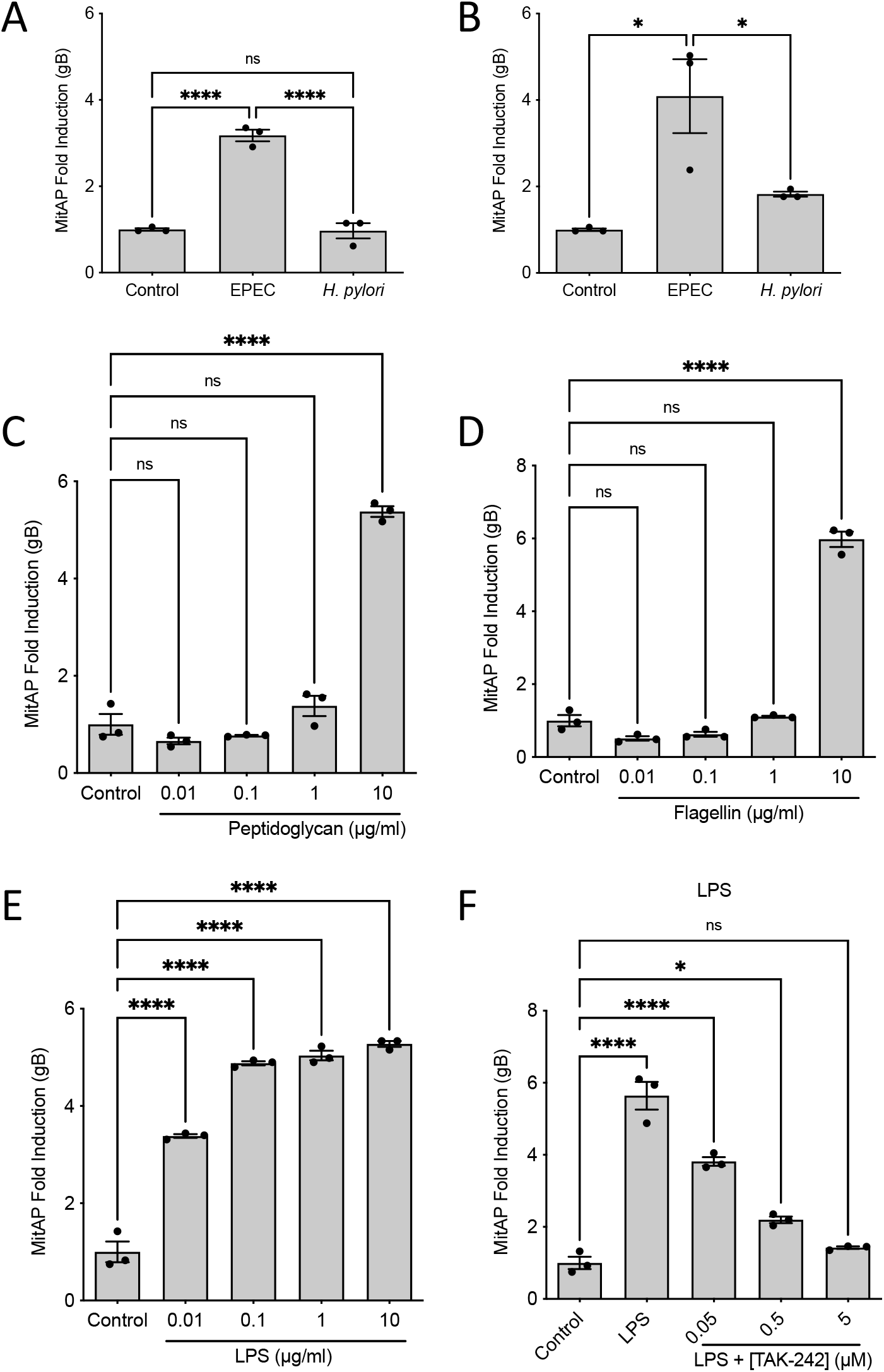
LPS activates MitAP through TLR4. **A**) Of two Gram-negative bacteria, only EPEC activates MitAP. **B**) Similar results are observed with heat-killed bacteria, ruling out the need for an active inhibition. **C** and **D**) Peptidoglycan and Flagellin activate MitAP only at a very high concentration. **E**) LPS activates MitAP at a low concentration. **F**) TAK242, a specific TLR4 inhibitor, inhibits the activation of MitAP by LPS.

### The TRAM/TRIF arm of TLR4 signaling activates MitAP

To evaluate the contribution of the two arms of the TLR4 pathway, we first treated RAW macrophages with a specific inhibitory peptide of Myd88^12^. While this peptide effectively inhibited the release of IL-6 in response to LPS, demonstrating its efficacy, it had no inhibitory effect on the activation of MitAP (Fig. 2A). Similar results were observed when the downstream effector NF-κB was inhibited with BAY11-7082 (Fig. 2B), ruling out a major contribution of the Myd88-dependent arm of TLR4 in MitAP. The second arm of TLR4 involving TRAM and TRIF is activated from endosomes following the internalization by endocytosis of TLR4 and its ligand ^13^. In that context, we observed that Dynasore, a molecule that inhibits endocytosis, strongly inhibited the release of IFN-β, demonstrating its effect on the inhibition of the Tram/TRIF pathway. Dynasore also strongly inhibited the induction of MitAP by LPS (Fig. 2C). Treatment with piceatannol, a molecule that targets more specifically the Syk kinase/CD14 complex required for TLR4 internalization^13^, also inhibited the release of IFN-β and MitAP very efficiently (Fig. 2D). To complement these data, we used a genetic approach and produced CD14 KO RAW cells (CD14^-/-^) (Fig. 2E). We observed that TLR4 was no longer internalized in response to LPS in these cells (Fig. 2F). As a result, the release of IFN-β was inhibited and MitAP was not induced by LPS (Fig. 2G). These data clearly showed that the TRAM/TRIF arm of TLR4 signaling is involved in MitAP activation in inflammatory conditions. The role of the TRAM/TRIF arm in MitAP suggested that the transcription factor IRF3 is a downstream effector required for the activation of this antigen presentation pathway. To determine if this is the case, we produced an IRF3 KO RAW cell line (IRF3^-/-^) using CRISPR-Cas9 (Fig. 2H). As expected, the release of IFN-β, RANTES and CXCL10, three molecules whose expression is regulated by IRF3, was strongly decreased in the IRF3-/-cells in response to LPS (results not shown). Surprisingly, the induction of MitAP was not affected in IRF3^-/-^ cells (Fig. 2I), suggesting that a bifurcation along the TRAM/TRIF arm, upstream of IRF3, is responsible for MitAP activation. A key molecular complex involved in the TRAM/TRIF arm upstream of IRF3 is the kinase IKKε and its partner, the TANK-binding kinase-1 (TBK1), which phosphorylates IRF3 and promotes its translocation to the nucleus ^11^. Interestingly, inhibition of the TBK1/IKKε complex in RAW cells with the specific inhibitor BX795, which effectively inhibiter IRF3 phosphorylation, resulted in a significant inhibition of MitAP (Fig. 2J). Thus, our data indicate, so far, that following the activation of TLR4 by LPS, a pathway involving the TRAM/TRIF arm and the TBK1/IKKε complex, distinct from IRF3 activation, is engaged to induce MitAP.

**Fig. 2:**
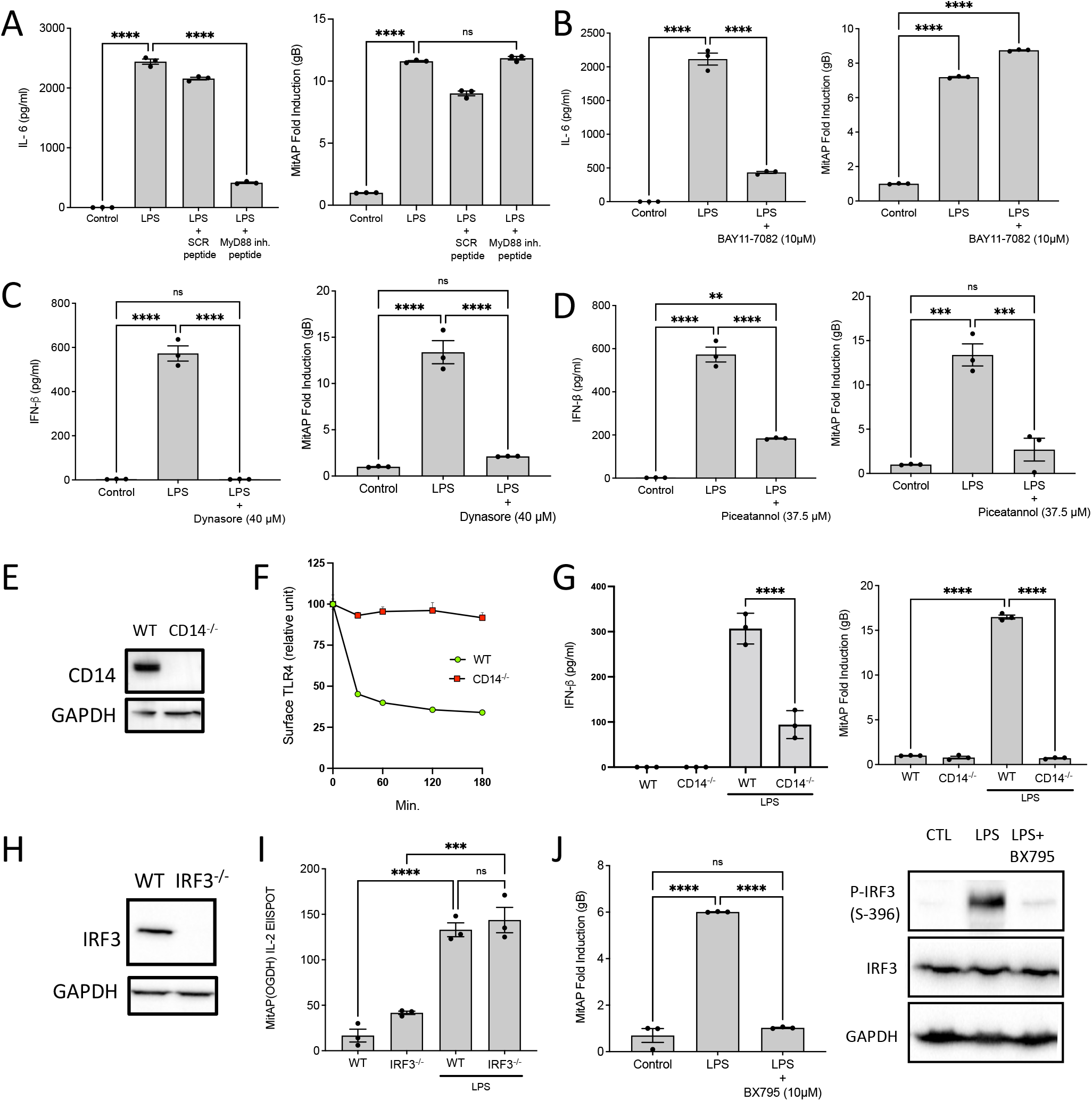
The TRAM/TRIF arm of TLR4 signaling activates MitAP. **A**) Inhibiting Myd88 with a specific peptide inhibits IL-6 release but not MitAP. **B**) Inhibiting NF-kB with Bay11-7082 inhibits IL-6 release (ELISA) but not MitAP. **C**) Inhibiting endocytosis with Dynasore inhibits both IFN-β release (ELISA) and MitAP. **D**) Inhibiting the internalization of CD14 with the Syk inhibitor Piceatannol inhibits both IFN-β release and MitAP. **E**) The loss of expression of CD14 in CD14^-/-^ RAW cells made with CRISPR/Cas9 was monitored by Western blotting. **F**) Flow cytometry indicates that the internalization of TLR4 from the cell surface is inhibited in response to LPS in CD14^-/-^ RAW cells. **G**) Both IFN-β and MitAP are inhibited in the CD14^-/-^ Raw cells. **H**) The loss of expression of IRF3 in IRF3^-/-^ RAW cells made with CRISPR/Cas9 was monitored by Western blotting. **I**) MitAP was not inhibited in response to LPS in IRF3^-/-^ RAW cells. **J**) Inhibition of TBK1 by BX795 inhibits MitAP (inhibition of P-IRF3 is shown to show the efficacy of BX795).

### STING is required to activate MitAP in inflammatory conditions

In APCs, the production of type-I IFN molecules such as IFN-β requires the stimulator of interferon genes (STING), which interacts with TBK1 to activate IRF3^14^. This interaction with TBK1 prompted us to test whether STING might play a role in the activation of MitAP, upstream of IRF3. First, we showed that the addition of cGAMP, an agonist of STING, to the culture medium was as potent as LPS at inducing MitAP (Fig. 3A). The activation of MitAP by cGAMP was not inhibited in IRF3^-/-^ cells, confirming that MitAP activation by STING occurred independently from IRF3 (Fig. 3B). We then produced STING KO RAW cells (STING^-/-^) to determine directly whether this protein is required for MitAP (Fig. 3C). We observed that MitAP was significantly inhibited in the STING^-/-^ following LPS treatment, compared to WT cells (Fig. 3D). It is well established that the cGAS-STING complex is activated by the presence of DNA in the cytoplasm which binds to cGAS, enabling the production of cGAMP which, in turn, binds to and activates STING (see ^15^). In WT RAW macrophages, we observed that LPS treatment led to the production of cGAMP (Fig. 3E), suggesting that DNA was released to the cytoplasm upon TLR4 activation. Previous studies provided evidence that mitochondrial DNA (mtDNA) is detected in the cytoplasm in response to LPS, activating cGAS-STING^16,17^. Interestingly, treatment of RAW macrophages with LPS led to the accumulation of mtDNA in small vesicular structures (Fig. 3F) reminiscent of mitochondria-derived vesicles (MDVs)^18^. Using qPCR, we observed that increasing levels of mtDNA coding for cytochrome B were detected in the cytoplasm in response to LPS (Fig. 3G). These data indicated that the release of DNA from mitochondria in response to LPS might be a mechanism by which STING activates MitAP.

**Fig. 3:**
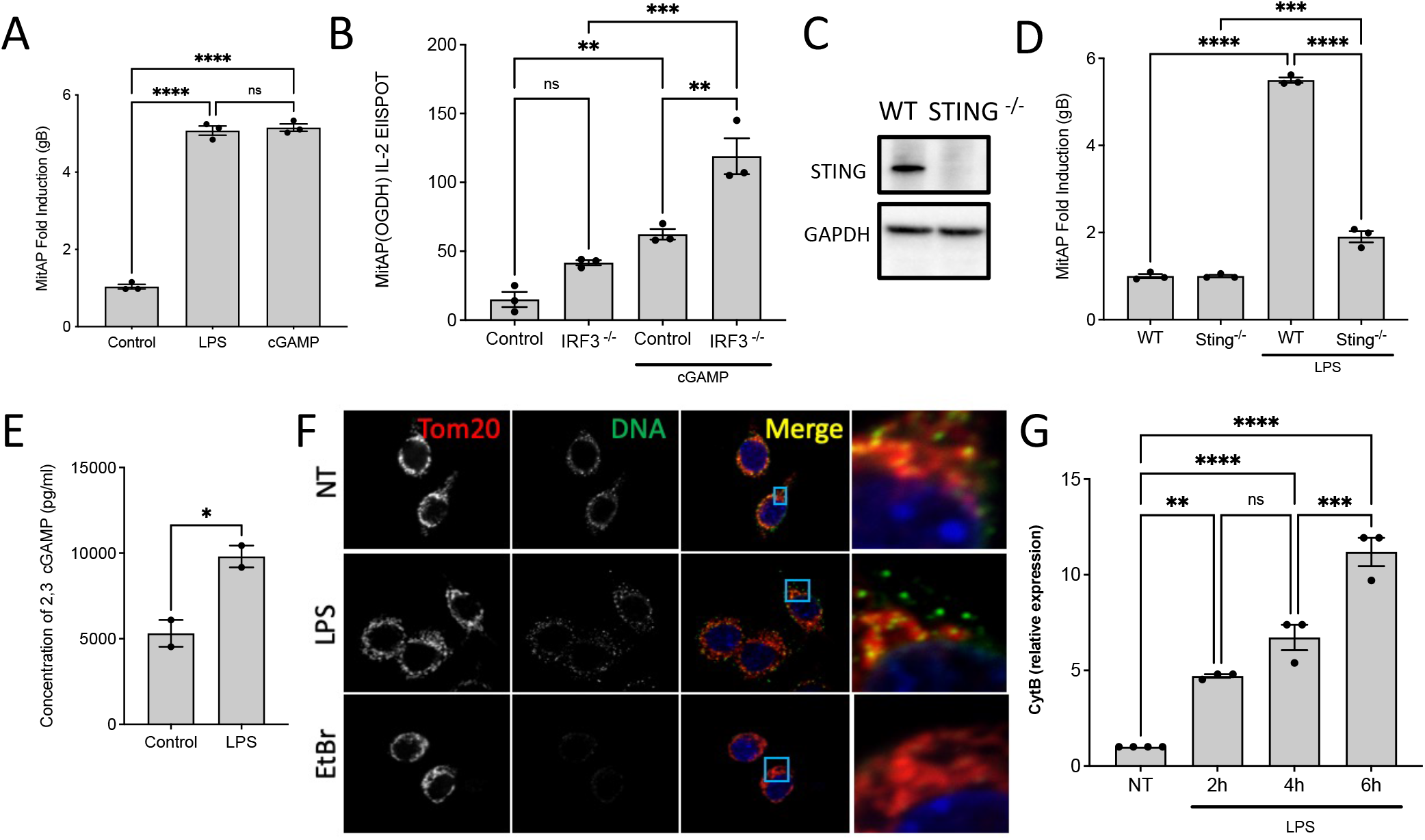
STING is required to activate MitAP in inflammatory conditions. **A**) The STING analogue cGAMP activates MitAP. **B**) IRF3 is not required for the activation of MitAP by cGAMP. **C**) The loss of expression of STING in STING^-/-^ RAW cells made with CRISPR/Cas9 is monitored by Western blotting. **D**) MitAP is strongly inhibited in STING^-/-^ cells in response to LPS. **E**) Treatment of RAW cells with LPS-induced cGAMP production. **F**) Immunofluorescence microscopy indicates that LPS treatment in RAW cells leads to the accumulation of mtDNA (DNA) in vesicular structures close to mitochondria (TOM20). The DNA signal is lost when mtDNA is depleted from mitochondria by ethidium bromide (EtBr) treatment (NT: Non-treated). **G**) LPS treatment in RAW cells increases the level of the mitochondrial CytB DNA detected by qPCR in the cytoplasm in a time-dependent manner.

While these data identified STING as a regulator of MitAP in inflammatory conditions, the finding that its two main downstream effectors, NF-κB and IRF3 ^19^, were not required for MitAP (see Figs. 2B and 2I) indicated that STING acts through a different pathway. A recent study reported that STING activation stimulates the Unfolded Protein Response (UPR) via a “UPR motif” within one of its cytoplasmic domains ^20^, highlighting a potential way by which STING could activate MitAP. The UPR is an adaptive signaling mechanism triggered by the accumulation of improperly folded proteins in the lumen of the endoplasmic reticulum (ER), aimed at restoring ER homeostasis ^21^. This stress response is transduced by three sensors: Protein Kinase R-like ER Kinase (PERK), Inositol Requiring Enzyme 1 (IRE1) and Activating Transcription Factor 6 (ATF6). As a first step to determine whether the UPR activates MitAP, we treated cells with two potent inducers of this pathway, Tunicamycin and Thapsigargin (TG) and observed that both molecules strongly activated MitAP in a dose-dependent manner (Supp. Fig. 2A). In contrast to LPS, activation of MitAP by TG was not inhibited by the TLR4 inhibitor TAK242 (Supp. Fig. 2B) and still occurred in cells lacking CD14 (Supp. Fig. 2C) or when TBK1 was inhibited with BX795 (Supp. Fig. 2D). These data indicated that the UPR triggered MitAP downstream of the TLR4 signaling events occurring in the cytoplasm. Finally, to further investigate whether the UPR played a role in MitAP, we treated cells with inhibitors of the three UPR sensors: Ceapin-A7 (ATF6); GSK2606414 (PERK); and Kira6 (IRE1) prior to LPS stimulation and observed a strong inhibition only with Kira6 (Supp. Fig. 2E), identifying IRE1 as a potent regulator of MitAP.

### STING engages MitAP through IRE1 activation

Next, we performed a series of experiments to assess whether STING and the UPR acted in coordination to activate the MitAP pathway. Using Western blot (WB) analyses, we showed that LPS treatment in RAW macrophages effectively led to the activation of the UPR, as shown by the production of XBP1s and ATF4, the downstream effectors of IRE1 and PERK respectively (Fig. 4A). Interestingly, in the absence of STING (STING ^-/-^), we observed that the production of XBP1s, but not ATF4, was strongly inhibited. ATF4 was, in fact, strongly upregulated (Fig. 4A). As expected, the secretion of the downstream effectors IL-6 and IFN-β was strongly inhibited in the STING^-/-^ cells (Fig. 4B). Concomitant, with the changes in the expression of the UPR effectors, MitAP activation in response to LPS was significantly inhibited in the STING^-/-^ cells (Fig. 4C). These data, along with the previous results with the UPR sensor inhibitors (Supp. Fig. 2E), supported the concept that STING is selectively regulating IRE1 to activate MitAP in inflammatory conditions, implying that STING is acting upstream of the UPR. This was confirmed by showing that MitAP activation was “rescued” in STING^-/-^ cells treated with TG (Fig. 4C). To test directly whether the “UPR motif” within STING was essential for the activation of the UPR and MitAP in response to LPS, we used CRISPR-Cas9 to edit the STING gene in RAW cells and introduced a double RA mutation in positions 330 and 333 (STING^RRAA^). These mutations (positions 331 and 334 in humans) were shown to abrogate the UPR motif function in human HEK293T cells^20^. As observed in the STING^-/-^ cells, the selective abrogation of the “UPR motif” function in STING^RRAA^ cells resulted in the specific inhibition of XBP1s production (Fig. 4D). Unlike the STING-/-, however, the release of IFN-β was not inhibited in STING^RRAA^ cells (Fig. 4E). Considering that MitAP was inhibited in the STING^RRAA^ cells (Fig. 4F), these results indicated that the mechanisms by which STING regulates the expression of IFN-β differs from those involved in the induction of IRE1 and MitAP. TG treatment in the STING^RRAA^ cells rescued both MitAP (Fig. 4F) and, to a large extent, the production of XBP1s (Fig. 4G), providing more evidence that this transcription factor is required for the activation of MitAP. To confirm the involvement of XBP1s in the MitAP pathway, we produced XBP1 KO RAW cells (XBP1^-/-^) and observed a significant decrease in the level of MitAP in response to LPS (Fig. 4H).

**Fig. 4:**
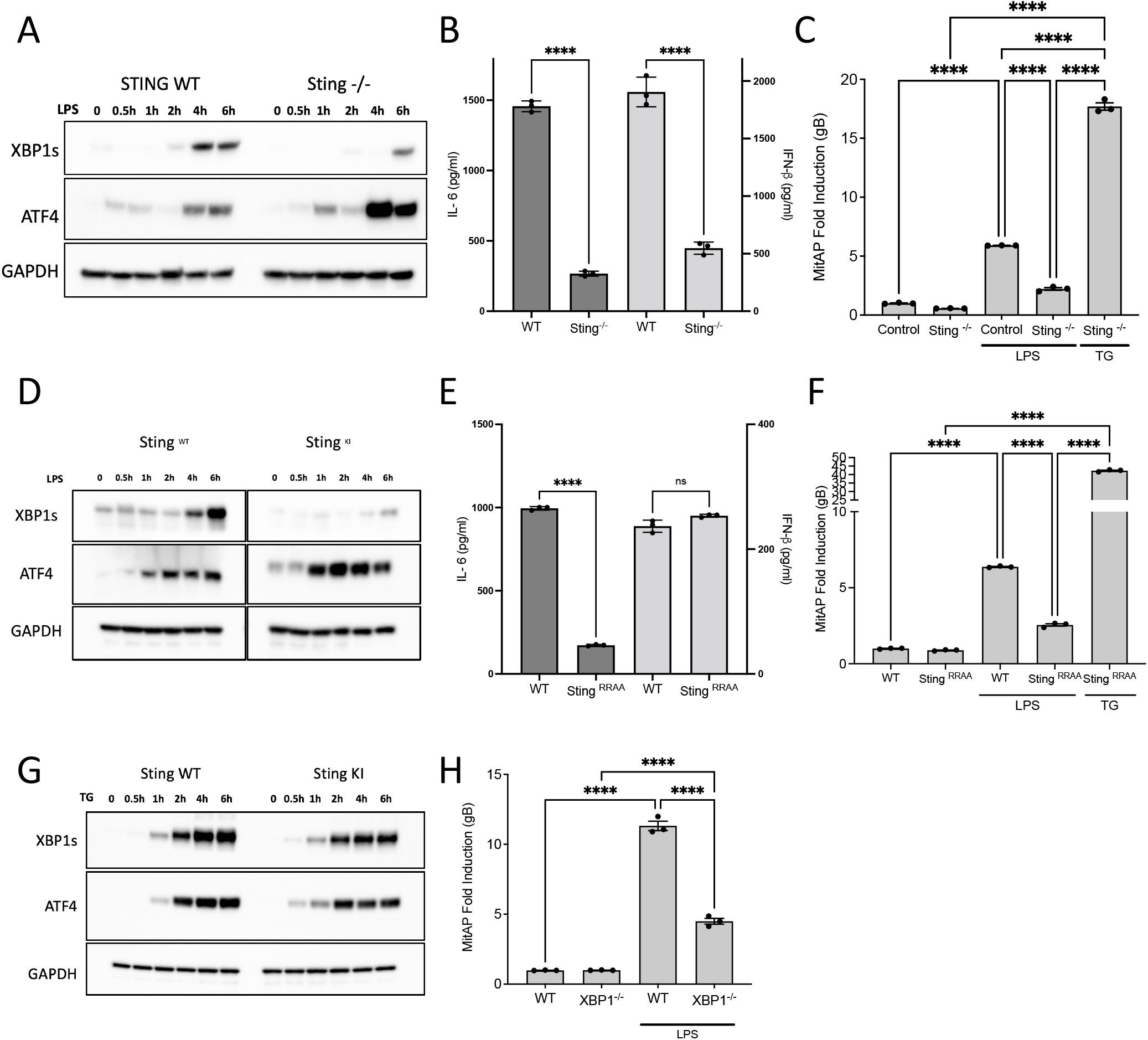
STING engages MitAP through UPR activation. **A**) Both XBP1s and ATF4 are produced in response to LPD in WT RAW cells. In contrast, the production of XBP1s is strongly inhibited in the STING^-/-^ cells. **B**) The secretion of both IL-6 and NF-κB is inhibited in STING^-/-^ cells in response to LPS. **C**) MitAP is inhibited in STING^-/-^ RAW cells in response to LPS but rescued by TG treatment. **D**) The production of XBP1 is also inhibited in STING^RRAA^ in response to LPS. **E**) STING^RRAA^ cells have a defect in the secretion of IL-6 but not IFN-β. **F**) A strong inhibition of MitAP in response to LPS is observed in STING^RRAA^ RAW cells, rescued by TG treatment. **G**) TG treatment rescues the expression of XBP1s protein and restores the level of ATF4 protein expression in the STING^RRAA^ RAW cells. **H**) MitAP is inhibited in response to LPS in XBP1KO RAW cells.

### LRRK2 and STING act in close vicinity to activate IRE1 and MitAP

Having shown previously that PINK1 and Parkin regulate MitAP ^7^, we wanted to investigate whether other PD-related proteins were involved in this pathway. The first thing we did was to look whether the secretion of pro-inflammatory cytokines was altered in LRRK2 KO RAW macrophages (LRRK2^-/-^) that we produced by CRISPR-Cas9. LRRK2 was selected following the report that this protein is highly expressed in immune cells, suggesting its involvement in key immune functions ^22^. We observed that the level of IL-6 and IFN-β secreted in response to LPS was strongly inhibited in the LRRK2^-/-^ cells (Fig. 5A). This prompted us to determine whether the loss of LRRK2 affected the TLR4-to-STING-to-IRE1 pathway responsible for the activation of MitAP in the inflammatory conditions described here. Interestingly, the loss of LRRK2 phenocopied the effects of the loss of STING (or the inactivation of its “UPR motif”) on the MitAP pathway. Indeed, MitAP was poorly induced by LPS in LRRK2^-/-^ RAW cells (Fig. 5B). This inhibition was also observed in bone marrow-derived macrophages isolated from LRRK2 KO mice (LRRK2^-/-^) (Fig. 5C). Activation of IRE1, monitored by the level of XBP1s expression, was also strongly inhibited in the RAW LRRK2^-/-^ cells, with an increase in ATF4 (Fig. 5D). Interestingly, as observed in the STING^RRAA^ cells, LRRK2^-/-^ cells responded well to TG treatment, with a restored expression of XBP1s (Fig. 5E). These data suggest that LRRK2 and STING act in close vicinity to activate IRE1 and XBP1s production along the MitAP pathway in inflammatory conditions.

**Fig. 5:**
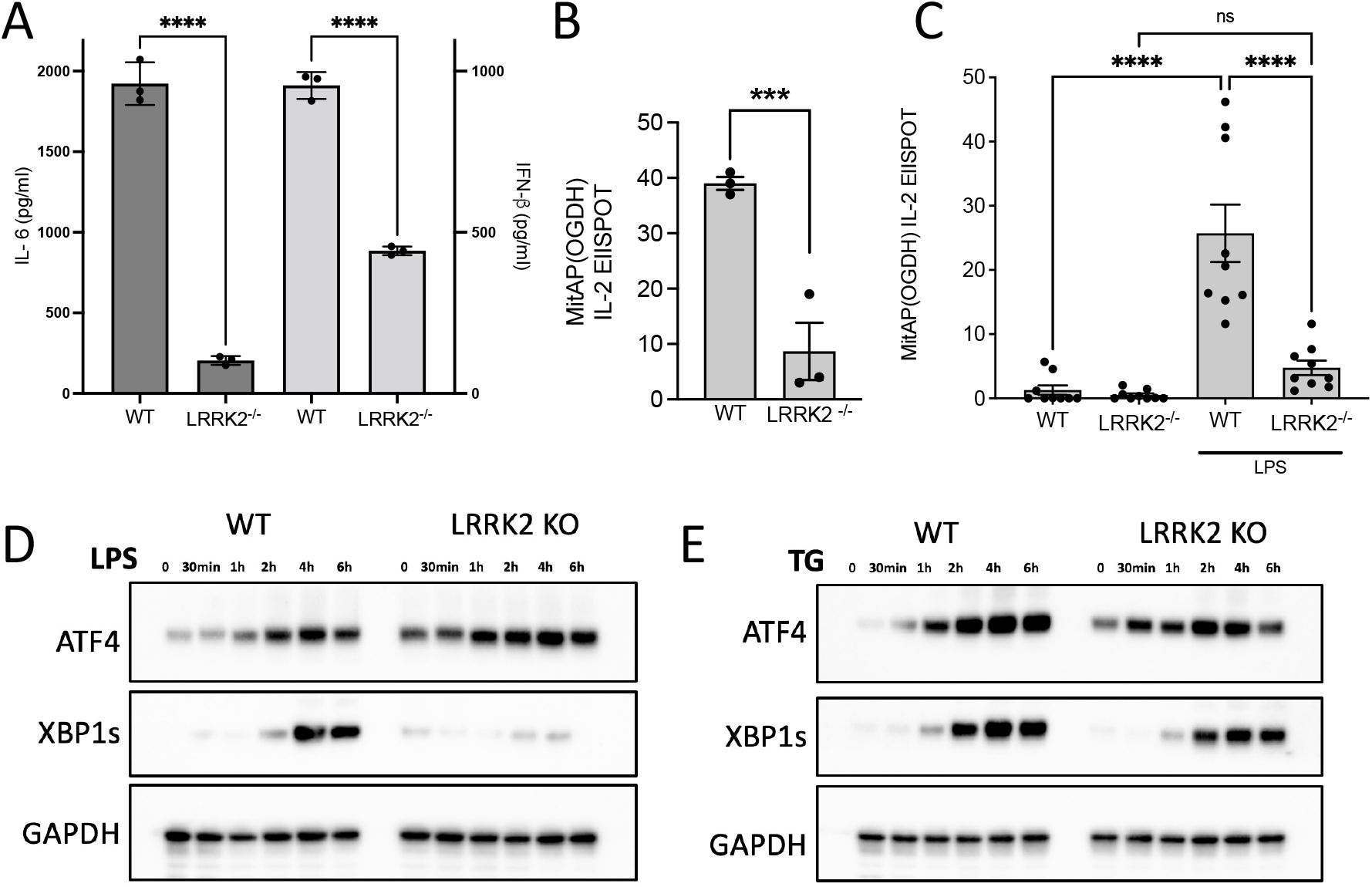
LRRK2 and STING act in close vicinity to activate IRE1 and MitAP. **A**) The secretion of both IL-6 and IFN-β (ELISA) was inhibited in LRRK2^-/-^ RAW cells in response to LPS **B**) MitAP is significantly inhibited in response to LPS in LRRK2^-/-^ RAW cells. **C**) A similar inhibition is observed in BMDMs isolated from LRRK2 KO mice. **D**) An inhibition in the level of expression of XBP1s protein in response to LPS, with an increase in ATF4, is observed in LRRK2^-/-^ RAW cells. **E**) TG treatment rescues the expression of XBP1s protein and restores the level of ATF4 protein expression in the LRRK2^-/-^ RAW cells.

## Discussion

With ageing, inflammation has been identified as a major risk factor for PD; “inflammageing” being presented as a persistent low-level state of inflammation fueled by chronically activated innate and adaptive immune cells in elderly people^3^. Immune senescence has been shown to increase the activation of autoimmune mechanisms^23^. A recent study identified autoreactive α-synuclein CD4^+^ T cells in PD patients^6^, suggesting that autoimmune mechanisms play a role in the development of PD. At the molecular cell level, we have shown that two PD-related proteins, PINK1 and Parkin, regulate the presentation of self-antigens derived from mitochondrial proteins during inflammation^7^. Indeed, activation of MitAP in *Pink1*^*-/-*^ mice following gut infection with Gram-negative bacteria led to the establishment of cytotoxic autoreactive T cells and the emergence of severe motor impairment, reversible by L-DOPA treatment^8^. Importantly, these studies identified PINK1 and Parkin as key regulators of the immune response during inflammation. The observation that MitAP also occurs in dopaminergic neurons, potentially allowing their recognition and attack by T cells, also suggested that autoimmune mechanisms play a role in the pathophysiological process leading to PD. In this context, we sought to characterize in molecular detail the mechanisms responsible for the presentation of antigens during inflammation and the role played by PD proteins in this process.

Macrophages are actively involved in both innate and adaptive immunity. Indeed, APCs are at the forefront of the inflammatory response as they participate in the initiation, maintenance and resolution of inflammation through the tight regulation of pro- and anti-inflammatory molecules secretion^24^. Macrophages also play a central role in the adaptive immune response by processing and presenting self or foreign antigens on MHC molecules at their surface to activate T cells at sites of inflammation. The molecular mechanisms enabling the activation of antigen presentation in response to inflammatory signals in APCs are still largely unknown. The pathway characterized here highlighted several novel aspects of how the transition from innate to adaptive immunity occurs in macrophages. First, we found that MitAP is driven by a complex pathway activated by multiple sensors along a TLR4-to-STING-to-IRE1 pathway. From the cell surface, the MitAP pathway is activated by the TRAM/TRIF arm of TLR4 signalling. Interestingly, IRF3, the main transcription factor activated by this arm of the pathway, was not required for the induction of MitAP by LPS. Indeed, we found that the signaling cascade initiated by the TRAM/TRIF arm bifurcates upstream of IRF3, at the level of TBK1, a kinase that interacts with and phosphorylates STING^25^. Our data showed that the cGAS-STING pathway turned out to be a second sensor for MitAP activation from the cytoplasm. We have shown that the product of cGAS, the cyclic dinucleotide cGAMP, is increased in response to LPS treatment, coincident with our observation that mtDNA is released into the cytosol. It is not yet clear how the TLR4-TRAM/TRIF arm may drive the release of mtDNA into the cytosol.

The apparent release of mtDNA in vesicular structures induced by LPS corroborates the observation made in a recent study showing that mtDNA reaches the cytoplasm from MDVs, where it activates STING^18^. This study further showed that the release of mtDNA to the cytoplasm required the cytosolic protein sorting nexin 9 (SNX9). Interestingly, we have shown previously that MitAP is also driven by the formation of MDVs in a SNX9-dependent manner ^7^, suggesting the involvement of a similar process of mtDNA release in the STING-dependent activation of MitAP described here.

Second, although further studies will determine whether mtDNA is required for MitAP, our data clearly identify STING as a key player in the activation of this presentation pathway during inflammation. The two prominent effectors downstream of STING, NFκB and IRF3 ^15^, turned out to play a minor role in MitAP. Instead, we found that the induction of STING by LPS in our pathway activates the UPR sensor IRE1, a step required for MitAP. Our data showing that XBP1s, but not ATF4, is inhibited in STING^RRAA^ RAW cells suggested that STING, through its “UPR motif”, may in fact selectively regulate the activity of IRE1. The specific involvement of IRE1 is supported by results showing that MitAP is not affected when cells are treated with PERK and ATF6 inhibitors, while the IRE1 inhibitor Kira6 completely inhibited MitAP. This goes along with a recent study showing that PERK does not play a major role in MHC-I antigen presentation^26^.

Abrogation of XBP1s expression in KO RAW cells, on the other hand, significantly inhibited MitAP. The regulation of the expression of a number of genes involved in antigen presentation on MHC-I molecules by the transcription factor XBP1s, such as Calnexin, Calreticulin and Tapasin, highlights how impairment of key upstream proteins affecting the activation of IRE1 may impair the immune system and promote autoimmune mechanisms.

Remarkably, we found that LRRK2 regulates the activation of the UPR in inflammatory conditions; impairment of this protein affecting both the level of expression of the XBP1s and ATF4 proteins, as well as the activation of MitAP. LRRK2 is thus the third PD-related protein, with PINK1 and Parkin ^7^, to be involved in the regulation of the adaptive immune response during inflammation. The potential role of LRRK2 in the immune system was highlighted by studies showing that this protein is highly expressed in immune cells and linked to immune-related diseases including leprosy and inflammatory bowel disease ^22^. It is of interest to note that the loss of LRRK2 in RAW cells phenocopies the alterations observed in both STING KO and STING^RRAA^ cells. In all of these cells, a strong inhibition of the expression of the IRE1-driven protein XBP1 is observed with a strong increase in the level of ATF4, potentially occurring as a compensatory mechanism. Interestingly, it was shown recently that virally-induced expression of ATF4 in the substantia nigra caused dopaminergic neuron loss in a rat model of PD overexpressing human α-synuclein^27^. Overactivation of STING increases neuroinflammation and causes DNs cell death^28^. In addition, the activation of the TBK1/STING axis during α-synucleinopathy was shown to mediate neuroinflammation and neurodegeneration^29^. The finding that LRRK2 and STING are closely acting to activate the UPR, along a pathway that regulates the inflammatory response, provides further mechanistic insights into the pathophysiological process that may lead to PD. Finally, in addition to the immune-related diseases mentioned above, LRRK2 was identified as a cancer predisposition gene in a recent study on the genomic landscape of familial glioma ^30^. All of these diseases have been linked to impairment of the UPR ^31-33^.

Our data are of particular interest when discussed in the context of the large number of genes associated with PD and its long prodromal period (it has been proposed that PD is initiated in peripheral organs, several years, often decades, before the onset of movement disorder) ^34^. The finding that LRRK2, in addition to PINK1 and Parkin ^7^, plays a role in regulating the engagement of antigen presentation in APCs during inflammation, positions for the first time three PD-related proteins along the same pathway. The need to tightly regulate this process to prevent autoimmune mechanisms is of further relevance to the long prodromal period of the disease. Indeed, we showed that MitAP activation during gut infection in *Pink1*^*-/-*^ mice is accompanied by the establishment of cytotoxic CD8^+^ T cells, detected in the brain of these animals. Unlike inflammation, which is a rapid (non-specific) immune response to stress, the activation of T cell populations enables a long-term “immune memory”, that can last for decades^35^. In a model where MitAP occurs first in APCs in the periphery and later in neurons, DNs would be “invisible” to the immune system until signals like neuroinflammation, associated with ageing ^3^, trigger MitAP in these cells rendering them susceptible to T cell attack. Antigen presentation in DNs was shown to promote T cell attack in vitro ^8,36^. Thus, targeting the MitAP pathway may allow the development of therapeutic approaches applicable at both early and late stages of the disease.

## Acknowledgements

The study is funded by the joint efforts of The Michael J. Fox Foundation for Parkinson’s Research (MJFF) and the Aligning Science Across Parkinson’s (ASAP) initiative. MJFF administers the grant ASAP 000525 on behalf of ASAP and itself. This work was also supported by CIHR grants to ES, SG, HM and MD. The authors wish to thank Dr. Auten Milnerwood and Camilla Tiefensee Ribeiro for the kind gift of *lrrk2-/-* mice.

## Materials and Methods

### Animals and cells

C57Bl/6J WT and LRRK2 KO mice were maintained according to the Canadian Council on Animal Care regulations. RAW 264.7 macrophage cell lines expressing H-2K^b^ and the glycoprotein B (gB) from HSV-1 in the mitochondria matrix was used as previously described ^7^. Cells were cultured in DMEM with 10% (v/v) fetal calf serum (FCS), penicillin (100 U ml^−1^) and streptomycin (100 μg ml^−1^). The β-galactosidase-inducible HSV gB/K^b^-restricted HSV-2.3.2E2 CD8^+^ T cell hybridoma (2E2) was provided by F. Carbone. The OGDH/L^d^ and OGDH/K^b^-restricted 2CZ CD8T^+^ cell hybridoma was provided by N. Shastri. Hybridomas were maintained in RPMI-1640 medium supplemented with 5% (v/v) FCS, glutamine (2 mM), penicillin (100 U ml^−1^) and streptomycin (100 μg ml^−1^).

### Reagents and Antibodies

Thapsigargin (T9033-1MG), Tunicamycin (<u>T7765-1MG</u>), ceapin-a7(SML2330-5MG), PERK Inhibitor I, GSK2606414(516535-5MG), RE1 Inhibitor IV, KIRA6 (532281), TAK-242(<u>614316-5MG</u>), Dynasore hydrate (D7693-5MG), Piceatannol (527948-1MG) were purchased from Sigma. BAY11-7082 (tlrl-b82), LPS-EB Ultrapure (TLRL-3PELPS), 2’3’-cGAMP (TLRL-NACGA23-5), PGN-EB (tlrl-pgneb), Fla-st (TLRL-STFLA), BX795 (tlrl-bx7) were from InvivoGen. MyD88 Inhibitor Peptide (NBP2-29328) was from R&D.

Antibodies against STING (13647S), CD14(93882S), IRF-3 (4302S), Phospho-IRF-3 (4947S), XBP-1s (12782S), ATF-4 (11815S) were purchased from Cell Signalling. Anti-GAPDH (MAB374) was from Sigma. Anti-Tom20 (ab115746) was from Abcam. Anti-DNA (61014) was from PROGEN.

### Bone marrow-derived macrophages (BMDMs) preparation

For the differentiation of BMDMs, sex-paired cohorts of male or female mice at 6 to 12 weeks of age were euthanized and bone marrow cells were harvested from the femurs and differentiated in DMEM + GlutaMAX-I (GIBCO) containing 10% FBS, 100 units/ml penicillin-streptomycin, 20 ng/ml recombinant mouse M-CSF (R&D/PeproTech) for 7 days. BMDM were then harvested and seeded on tissue culture plates 1 day before stimulation and maintained in complete DMEM media.

### Western Blotting

Cells were lysed in Laemmli buffer and the protein content quantified using EZQ™ protein quantification kit (Thermo Fisher). Equal protein amounts were loaded and separated on 4%– 15% pre-cast gels (BioRad) and transferred using a Trans-Blot turbo system (BioRad).

Membranes were blocked either in 5% milk TBS or 5%BSA TBS for 30 min at room temperature and incubated with the primary antibody diluted in 5% BSA-TBS overnight at 4 °C. The following day, the membrane was washed 3 times in Tris-buffered saline -Tween(20)(TBS-T), incubated with the secondary antibody (diluted in TBS-T plus 5% milk or 5% BSA) for 1h at room temperature and washed 3 times in TBS-T. The membrane was then developed in Clarity Western ECL substrate (BioRad) and visualized using the ChemiDoc imaging system (BioRad).

### ELISA

Collected supernatants were assayed for IL-6 and IFN-β using mouse DuoSet kits (R&D Systems). Samples were diluted in PBS and incubated in a 96-well plate pre-coated with either IL-6 or IFN-β capture antibodies for 2h at room temperature. Plates were then washed three times with PBS plus 0.05% Tween20 and incubated with the respective detection antibody for 2h at room temperature followed by three washes. Plates were finally incubated with streptavidin–HRP in the dark for 20 min, washed three times and developed with a 1:1 ratio of hydrogen peroxide and tetramethylbenzidine (Thermo Fisher). The optical density of the plates was read at 450 nm using a Varioskan microplate reader (Thermo Fisher). 2’’3’’-cGAMP ELISA was done using the 2’’3’’-cGAMP ELISA Kit (Cayman Chemical Co.) according to the manufacturer’s instructions.

### MHC class I antigen-presentation assay

One day prior to treatment, RAW cells were plated in a 24-well plate at a confluency of (1×10^6^/well). The following day, cells were either infected with EPEC at an M.O.I. of 1 or treated with indicated inducers for 6h. In the case of the tested inhibitors, cells were treated with the inhibitor for 2h prior to stimulation with LPS. Cells were then transferred into a 96-well plate 1h prior to fixation. Cells were fixed with 1% paraformaldehyde (PFA) for 10 min at room temperature, washed 5 times with wash media (RPMI, 10% FBS and 0.1 M glycine) and incubated with 1 × 10^5^ 2E2 T cell hybridoma for 16 h. β-galactosidase activity was then measured at 595 nm after the addition of CPRG substrate using a Varioskan microplate reader (Thermo Fisher). For OGDH antigen presentation in RAW cells or BMDMs expressing the H2-L^d^ allele, the 2CZ T cell hybridoma was used and the IL-2 secreted upon its activation was quantified using an ELISPOT assay (Mabtech) as previously described ^7^.

### Immunofluorescence

RAW cells were fixed with 5% PFA for 15 min at 37 °C. The PFA was then quenched with 50 mM NH_4_Cl in PBS for 10 min at room temperature. Cells were permeabilized with 0.1% Triton X-100 in PBS (v/v) for 10 min at room temperature then blocked with blocking buffer (5% FBS in PBS) for 10 min at room temperature. Cells were incubated with the indicated primary antibodies for 16 h at 4°C. After three washes with PBS, cells were incubated with the appropriate secondary antibodies (anti-rabbit-A488 Life Technologies, 1:1000; anti-mouse-A568 Life Technologies, 1:1000) for 1 h. After being washed and mounted with Prolong™ Antifade (Invitrogen), the slides were examined with a Zeiss LSM 780 laser scanning confocal microscope. For depletion of mtDNA, cells were treated with EtBr (200ng/ml) for 3 days then treated or not with LPS for 6 h and then fixed and immunostained as described above.

### Crispr-Cas9 gene editing

The gRNAs for the generation of knock-out (KO) and knock-in (KI) RAW cells were synthesized using GeneArt precision gRNA synthesis kits (Thermo Fisher) according to the manufacturer’s instructions. Prior to making gRNAs, 34-nucleotide forward and reverse target DNA oligonucleotides were designed using the CRISPR search and design tool (Thermo Fisher) and synthesized. The gRNA DNA templates were then PCR assembled and gRNAs were synthesized by *in vitro* transcription. Then gRNAs were purified, and their concentrations assessed. Prior to electroporation, Raw cells were scraped, washed with PBS and counted. 1 × 10^5^ cells were then electroporated with 2μg Cas9 protein and 400 ng gRNA in 10 μl of resuspension buffer R (and 50 pmol of donor HDR templates in case of KI) using Neon transfection system (Thermo Fisher) according to the manufacturer’s instructions. After electroporation, cells were immediately added into prewarmed 1 ml growth medium in a well of a 12-well plate and cultured for 4 days. Cells were counted and serially diluted to 4 cells/ml. 200 μl of cell suspension was dispensed in each well of a 96-well plate. Plates were then incubated at 37°C in a 5% CO_2_ incubator. Genomic DNA was isolated from single clones. The gRNA target region was amplified by PCR using AmpliTaq Gold 360 master mix (Thermo Fisher). PCR amplicons were sequenced using standard Sanger sequencing. KO cells were also confirmed by Western blotting.

### RT-qPCR

RNA was extracted using Aurum™ Total RNA Mini Kit (Biorad). Isolated RNA was then reverse transcribed with SuperScript™ VILO™ cDNA Synthesis Kit (Thermo Fisher). The generated cDNA was used for qPCR using Taqman probes (Thermo Fisher) and results were analyzed using the comparative ΔCt method.

### Quantification of mtDNA by qPCR

RAW cells (4×10^6^) were plated in 6 well plate and stimulated with LPS for the indicated time point. Then, cytosolic fraction was collected and mtDNA was isolated and quantified and previously described ^37^.

### Flow cytometry

RAW cells (0.5×10^6^) were stimulated with LPS for the indicated time points. Cells were then washed with 1 ml cold PBS. Live/dead cells were separated by ZombieAqua (BioLegend, #423102) and stained (30 min at 4°C) with PE anti-TLR4 (Biolegend; clone Sa15-21; 145404) in blocking buffer (PBS with 2% rat serum). The stained cells were then washed with 1 ml cold PBS and resuspended in 200 ml PBS. Staining of the surface receptors was analyzed with BD FACSCanto II. The mean fluorescence intensity (MFI) of TLR4 from unstimulated or LPS-stimulated cells was recorded. The percentage of surface receptor staining at the indicated time points, which is the ratio of the MFI values measured from the stimulated cells to those measured from the unstimulated cells, was plotted to reflect the efficiency of receptor endocytosis.

### Statistical analysis

The results shown in this study are representative of at least three independent experiments. Student’s-t-test and one-way ANOVA with Dunnett’s post-test were performed using GraphPad Prism 9. P-values below 0.05 were considered statistically significant.

## Figure legends

**Supp. Fig. 1:**
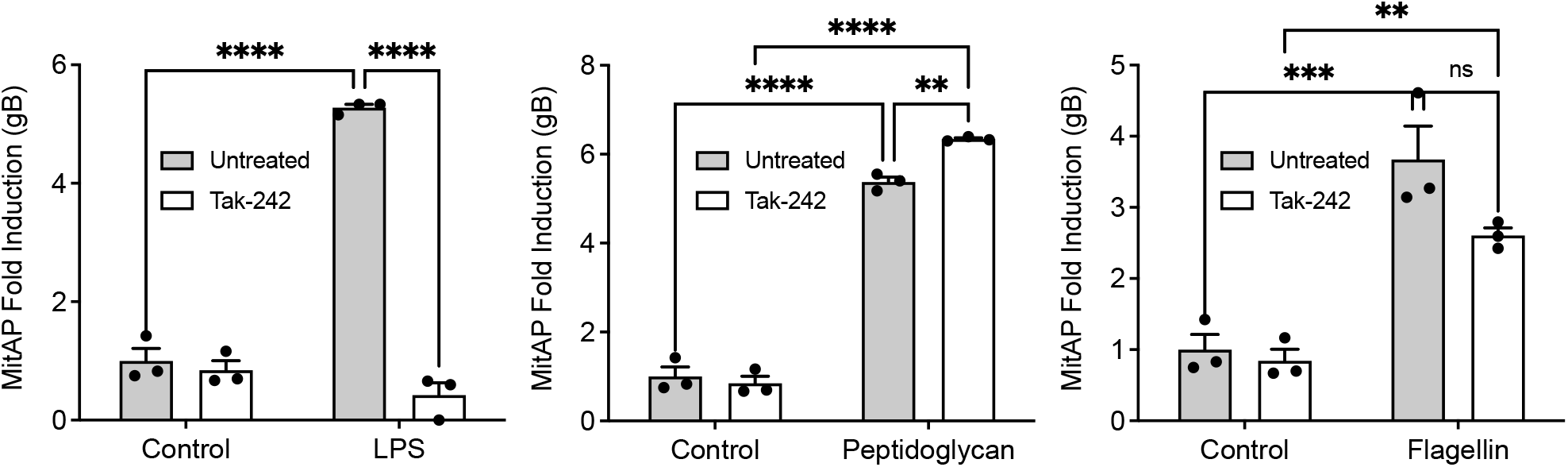
LPS activates MitAP through TLR4. The activation of MitAP by LPS, but not by a high concentration of peptidoglycan and flagellin, is inhibited by a pre-treatment with the specific TLR4 inhibitor TAK242.

**Supp. Fig. 2:**
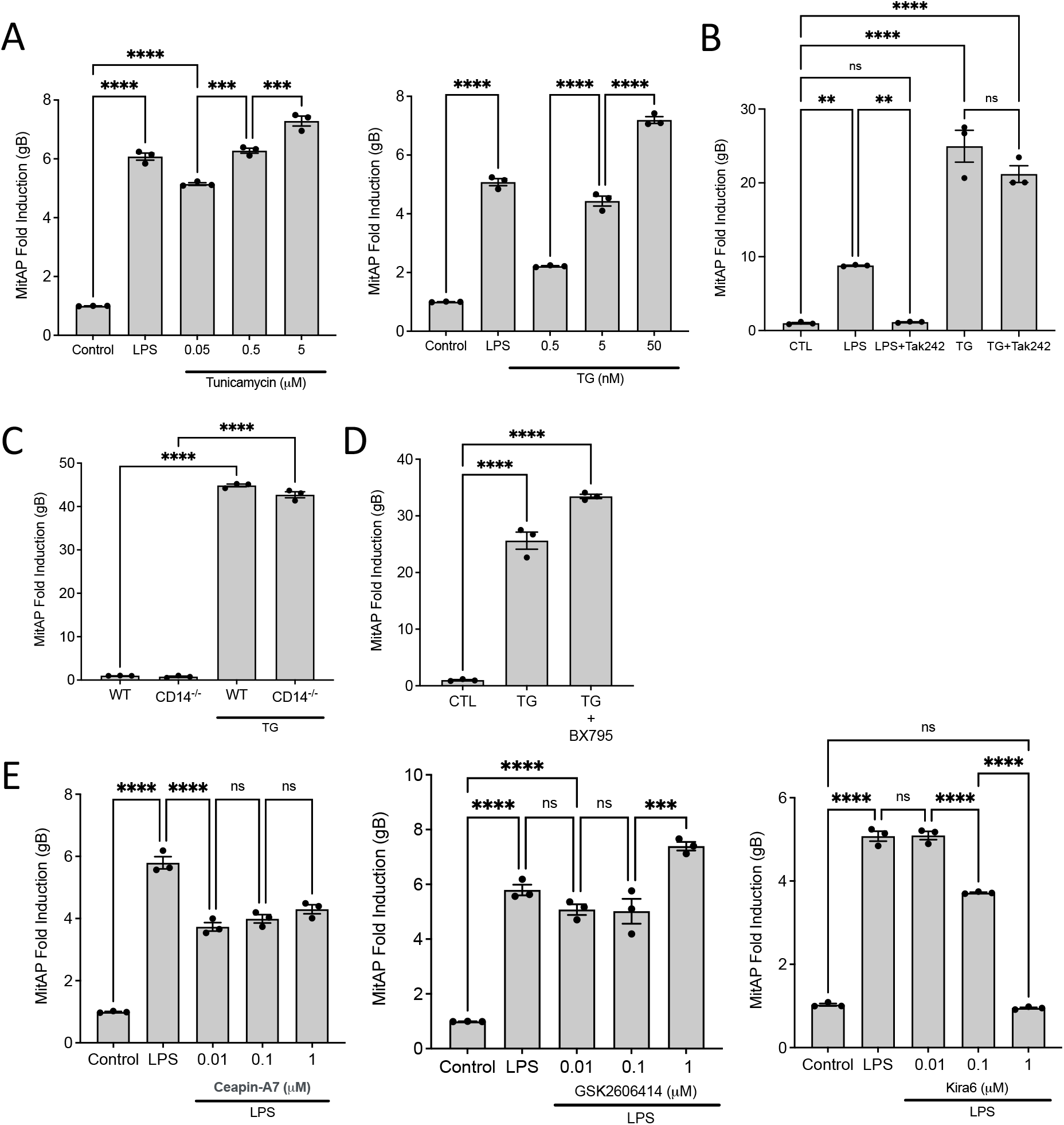
The UPR activates MitAP downstream of STING. **A**) Tunicamycin and Thapsigargin (TG) treatment induce MitAP in a dose-dependent way in WT RAW macrophages. **B**) Activation of MitAP by TG is not inhibited by TAK242. **C**) Activation of MitAP by TG is not inhibited in the CD14^-/-^ RAW cells. **D**) Activation of MitAP by TG is not inhibited by BX795 (TBK1 inhibitor). **E**) Treatment with the inhibitors Ceapin-A7 (ATF6) and GSK2606414 (PERK) has no effect on the level of MitAP induced by LPS. In contrast, treatment with the inhibitor Kira6 (IRE1) inhibits MitAP in a dose-dependent manner.

## References

1. Nalls, M.A., Blauwendraat, C., Vallerga, C.L., Heilbron, K., Bandres-Ciga, S., Chang, D., Tan, M., Kia, D.A., Noyce, A.J., Xue, A., et al. (2019). Identification of novel risk loci, causal insights, and heritable risk for Parkinson’s disease: a meta-analysis of genome-wide association studies. Lancet Neurol 18, 1091–1102. 10.1016/S1474-4422(19)30320-5.

2. Mogi, M., Harada, M., Kondo, T., Riederer, P., Inagaki, H., Minami, M., and Nagatsu, T. (1994). Interleukin-1 beta, interleukin-6, epidermal growth factor and transforming growth factor-alpha are elevated in the brain from parkinsonian patients. Neurosci Lett 180, 147–150. 10.1016/0304-3940(94)90508-8.

3. Tansey, M.G., Wallings, R.L., Houser, M.C., Herrick, M.K., Keating, C.E., and Joers, V. (2022). Inflammation and immune dysfunction in Parkinson disease. Nat Rev Immunol 22, 657–673. 10.1038/s41577-022-00684-6.

4. McGeer, P.L., Itagaki, S., Akiyama, H., and McGeer, E.G. (1988). Rate of cell death in parkinsonism indicates active neuropathological process. Ann Neurol 24, 574–576. 10.1002/ana.410240415.

5. Brochard, V., Combadiere, B., Prigent, A., Laouar, Y., Perrin, A., Beray-Berthat, V., Bonduelle, O., Alvarez-Fischer, D., Callebert, J., Launay, J.M., et al. (2009). Infiltration of CD4+ lymphocytes into the brain contributes to neurodegeneration in a mouse model of Parkinson disease. J Clin Invest 119, 182–192. 10.1172/JCI36470.

6. Sulzer, D., Alcalay, R.N., Garretti, F., Cote, L., Kanter, E., Agin-Liebes, J., Liong, C., McMurtrey, C., Hildebrand, W.H., Mao, X., et al. (2017). T cells from patients with Parkinson’s disease recognize alpha-synuclein peptides. Nature 546, 656–661. 10.1038/nature22815.

7. Matheoud, D., Sugiura, A., Bellemare-Pelletier, A., Laplante, A., Rondeau, C., Chemali, M., Fazel, A., Bergeron, J.J., Trudeau, L.E., Burelle, Y., et al. (2016). Parkinson’s Disease-Related Proteins PINK1 and Parkin Repress Mitochondrial Antigen Presentation. Cell 166, 314–327. 10.1016/j.cell.2016.05.039.

8. Matheoud, D., Cannon, T., Voisin, A., Penttinen, A.M., Ramet, L., Fahmy, A.M., Ducrot, C., Laplante, A., Bourque, M.J., Zhu, L., et al. (2019). Intestinal infection triggers Parkinson’s disease-like symptoms in Pink1(-/-) mice. Nature 571, 565–569. 10.1038/s41586-019-1405-y.

9. Lu, Y.C., Yeh, W.C., and Ohashi, P.S. (2008). LPS/TLR4 signal transduction pathway. Cytokine 42, 145–151. 10.1016/j.cyto.2008.01.006.

10. Yokota, S., Ohnishi, T., Muroi, M., Tanamoto, K., Fujii, N., and Amano, K. (2007). Highly-purified Helicobacter pylori LPS preparations induce weak inflammatory reactions and utilize Toll-like receptor 2 complex but not Toll-like receptor 4 complex. FEMS Immunol Med Microbiol 51, 140–148. 10.1111/j.1574-695X.2007.00288.x.

11. Kawasaki, T., and Kawai, T. (2014). Toll-like receptor signaling pathways. Front Immunol 5, 461. 10.3389/fimmu.2014.00461.

12. Jacobovitz, M.R., Rupp, S., Voss, P.A., Maegele, I., Gornik, S.G., and Guse, A. (2021). Dinoflagellate symbionts escape vomocytosis by host cell immune suppression. Nat Microbiol 6, 769–782. 10.1038/s41564-021-00897-w.

13. Zanoni, I., Ostuni, R., Marek, L.R., Barresi, S., Barbalat, R., Barton, G.M., Granucci, F., and Kagan, J.C. (2011). CD14 controls the LPS-induced endocytosis of Toll-like receptor 4. Cell 147, 868–880. 10.1016/j.cell.2011.09.051.

14. Ishikawa, H., Ma, Z., and Barber, G.N. (2009). STING regulates intracellular DNA-mediated, type I interferon-dependent innate immunity. Nature 461, 788–792. 10.1038/nature08476.

15. Barber, G.N. (2015). STING: infection, inflammation and cancer. Nat Rev Immunol 15, 760–770. 10.1038/nri3921.

16. Zhan, X., Cui, R., Geng, X., Li, J., Zhou, Y., He, L., Cao, C., Zhang, C., Chen, Z., and Ying, S. (2021). LPS-induced mitochondrial DNA synthesis and release facilitate RAD50-dependent acute lung injury. Signal Transduct Target Ther 6, 103. 10.1038/s41392-021-00494-7.

17. Zhou, L., Zhang, Y.F., Yang, F.H., Mao, H.Q., Chen, Z., and Zhang, L. (2021). Mitochondrial DNA leakage induces odontoblast inflammation via the cGAS-STING pathway. Cell Commun Signal 19, 58. 10.1186/s12964-021-00738-7.

18. Zecchini, V., Paupe, V., Herranz-Montoya, I., Janssen, J., Wortel, I.M.N., Morris, J.L., Ferguson, A., Chowdury, S.R., Segarra-Mondejar, M., Costa, A.S.H., et al. (2023). Fumarate induces vesicular release of mtDNA to drive innate immunity. Nature 615, 499–506. 10.1038/s41586-023-05770-w.

19. Yum, S., Li, M., Fang, Y., and Chen, Z.J. (2021). TBK1 recruitment to STING activates both IRF3 and NF-kappaB that mediate immune defense against tumors and viral infections. Proc Natl Acad Sci U S A 118. 10.1073/pnas.2100225118.

20. Wu, J., Chen, Y.J., Dobbs, N., Sakai, T., Liou, J., Miner, J.J., and Yan, N. (2019). STING-mediated disruption of calcium homeostasis chronically activates ER stress and primes T cell death. J Exp Med 216, 867–883. 10.1084/jem.20182192.

21. Almanza, A., Carlesso, A., Chintha, C., Creedican, S., Doultsinos, D., Leuzzi, B., Luis, A., McCarthy, N., Montibeller, L., More, S., et al. (2019). Endoplasmic reticulum stress signalling - from basic mechanisms to clinical applications. FEBS J 286, 241–278. 10.1111/febs.14608.

22. Wallings, R.L., Herrick, M.K., and Tansey, M.G. (2020). LRRK2 at the Interface Between Peripheral and Central Immune Function in Parkinson’s. Front Neurosci 14, 443. 10.3389/fnins.2020.00443.

23. Goronzy, J.J., and Weyand, C.M. (2013). Understanding immunosenescence to improve responses to vaccines. Nat Immunol 14, 428–436. 10.1038/ni.2588.

24. Fujiwara, N., and Kobayashi, K. (2005). Macrophages in inflammation. Curr Drug Targets Inflamm Allergy 4, 281–286. 10.2174/1568010054022024.

25. Zhang, C., Shang, G., Gui, X., Zhang, X., Bai, X.C., and Chen, Z.J. (2019). Structural basis of STING binding with and phosphorylation by TBK1. Nature 567, 394–398. 10.1038/s41586-019-1000-2.

26. Nagamine, B.S., Godil, J., and Dolan, B.P. (2021). The Unfolded Protein Response Reveals eIF2alpha Phosphorylation as a Critical Factor for Direct MHC Class I Antigen Presentation. Immunohorizons 5, 135–146. 10.4049/immunohorizons.2100012.

27. Gully, J.C., Sergeyev, V.G., Bhootada, Y., Mendez-Gomez, H., Meyers, C.A., Zolotukhin, S., Gorbatyuk, M.S., and Gorbatyuk, O.S. (2016). Up-regulation of activating transcription factor 4 induces severe loss of dopamine nigral neurons in a rat model of Parkinson’s disease. Neurosci Lett 627, 36–41. 10.1016/j.neulet.2016.05.039.

28. Szego, E.M., Malz, L., Bernhardt, N., Rosen-Wolff, A., Falkenburger, B.H., and Luksch, H. (2022). Constitutively active STING causes neuroinflammation and degeneration of dopaminergic neurons in mice. Elife 11. 10.7554/eLife.81943.

29. Hinkle, J.T., Patel, J., Panicker, N., Karuppagounder, S.S., Biswas, D., Belingon, B., Chen, R., Brahmachari, S., Pletnikova, O., Troncoso, J.C., et al. (2022). STING mediates neurodegeneration and neuroinflammation in nigrostriatal alpha-synucleinopathy. Proc Natl Acad Sci U S A 119, e2118819119. 10.1073/pnas.2118819119.

30. Choi, D.J., Armstrong, G., Lozzi, B., Vijayaraghavan, P., Plon, S.E., Wong, T.C., Boerwinkle, E., Muzny, D.M., Chen, H.C., Gibbs, R.A., et al. (2023). The genomic landscape of familial glioma. Sci Adv 9, eade2675. 10.1126/sciadv.ade2675.

31. Kaser, A., and Blumberg, R.S. (2013). Introduction: the unfolded protein response’s role in disease pathophysiology. Semin Immunopathol 35, 255–257. 10.1007/s00281-013-0379-3.

32. Obacz, J., Avril, T., Le Reste, P.J., Urra, H., Quillien, V., Hetz, C., and Chevet, E. (2017). Endoplasmic reticulum proteostasis in glioblastoma-From molecular mechanisms to therapeutic perspectives. Sci Signal 10. 10.1126/scisignal.aal2323.

33. Kumar, N., Khan, N., Cleveland, D., and Geiger, J.D. (2021). A common approach for fighting tuberculosis and leprosy: controlling endoplasmic reticulum stress in myeloid-derived suppressor cells. Immunotherapy 13, 1555–1563. 10.2217/imt-2021-0197.

34. Berg, D., Borghammer, P., Fereshtehnejad, S.M., Heinzel, S., Horsager, J., Schaeffer, E., and Postuma, R.B. (2021). Prodromal Parkinson disease subtypes - key to understanding heterogeneity. Nat Rev Neurol 17, 349–361. 10.1038/s41582-021-00486-9.

35. Kumar, B.V., Connors, T.J., and Farber, D.L. (2018). Human T Cell Development, Localization, and Function throughout Life. Immunity 48, 202–213. 10.1016/j.immuni.2018.01.007.

36. Cebrian, C., Zucca, F.A., Mauri, P., Steinbeck, J.A., Studer, L., Scherzer, C.R., Kanter, E., Budhu, S., Mandelbaum, J., Vonsattel, J.P., et al. (2014). MHC-I expression renders catecholaminergic neurons susceptible to T-cell-mediated degeneration. Nat Commun 5, 3633. 10.1038/ncomms4633.

37. Bronner, D.N., and O’Riordan, M.X. (2016). Measurement of Mitochondrial DNA Release in Response to ER Stress. Bio Protoc 6. 10.21769/BioProtoc.1839.

